# From Hot Water to Dry Dirt: Microbes Use Cytochrome ‘Nanowires’ of Similar Conductivity but Different Structure

**DOI:** 10.1101/2023.06.12.544705

**Authors:** Matthew J. Guberman-Pfeffer

## Abstract

Micron-scale electron transfer through polymeric cytochrome ‘nanowires’ powers prokaryotic life from hydrothermal vents to terrestrial soils in ways not fully understood. How much structural diversity optimizes electrical conductivity for survival in these different habitats is challenging to assess experimentally. Herein, physiologically relevant redox conduction is computationally assessed in cytochrome filaments from *Geobacter sulfurreducens* (OmcE, OmcS, and OmcZ), *Pyrobaculum calidifontis* (A3MW92), and *Archaeoglobus veneficus* (F2KMU8). A newly implemented Python program, BioDC, is used and validated against redox currents predicted from considerably more expensive molecular dynamics and quantum mechanical/molecular mechanical calculations. BioDC uses the heme solvent accessibility, stacking geometry, and redox-linked change in electrostatic energy to estimate electron transfer energetics. Leveraging this efficiency, structurally diverse cytochrome ‘nanowires’ from different organisms are shown to have similar redox conductivities. A functionally robust heme chain ‘packaged’ in habitat-customized proteins is proposed to be a general evolutionary design principle for cytochrome ‘nanowires’ widely distributed among prokaryotes.

**TOC Graphic:** 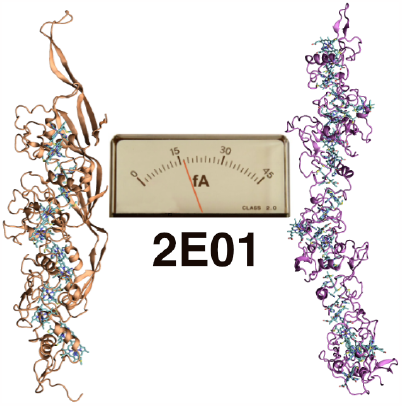

---

Microorganisms ranging from hyperthermophilic archaea to mesophilic proteobacteria extracellularly discharge electrons from hetero- or chemolithoautotrophic growth via filaments of multi-heme cytochromes.^1, 2^ These cytochrome ‘nanowires’ carry electrons across microbe-microbe^3^ and microbe-mineral^4, 5^ interfaces as the terminal step in an ancient^1^ respiratory strategy of bio- geochemical^4, 5^ and technological^6^ significance.

Structural determinations have rapidly increased in pace,^2, 7-11^ while electrical characterizations remain difficult,^2^ limited,^7, 12^ physiologically irrelevant,^2, 13-15^ and theoretically intringing.^13, 16-18^ Conducting-probe atomic force microscopy (CP-AFM) experiments indicate that two cytochrome ‘nanowires’ from the same proteobacterium show a 10^3^-fold variation in electrical conductivity when directly drop-casted on Au electrodes, dried, and compressed under nNs of force.^12^ Does this finding reflect how much evolution has optimized conductivity *under physiological conditions* through diversification of protein structure? Is even greater tunability realized when newly reported cytochrome ‘nanowires’ from hetero- and chemolithoautotrophic, hyperthermophilic archaea^2^ are considered? These fundamental questions, which are at the limits of experimental techniques, are addressed in the present Letter with the implementation of the theory for physiologically relevant^13, 19^ redox conduction^14, 20^ in the BioDC Python program, and its application to all presently known cytochrome ‘nanowires.’

The known filaments (Figure 1) are high-aspect ratio (>500 nm × <5 nm) helical homopolymers of outer-membrane cytochrome (Omc) types E (PDB ID 7TF7),^9^ S (PDB 6EF8 and 6NEF),^7, 8^ and Z (PDB 8D9M and 7LQ5)^10, 11^ from *Geobacter sulfurreducens*, A3MW92 from *Pyrobaculum calidifontis* (PDB 8E5F),^2^ and F2KMU8 from *Archaeoglobus veneficus* (PDB 8E5G).^2^ These proteins are tetra-, hexa-, and octa-heme cytochromes that follow a similar architectural blueprint: A chain of (typically) alternating parallel-displaced and perpendicularly stacked pairs of bis-histidine-ligated *c*-type heme cofactors are encased by a mostly (>50%) disordered protein sheath. However, the proteins show important differences in α-helical (13 – 29%) and β-sheet (3 – 18%) content, as well as the number (0, 1, or 3) of His residues coordinated to hemes in adjacent subunits of the filament. All these structures, including the models deposited by different labs for OmcS and OmcZ, are computationally characterized and compared in the present Letter for redox conductivity.

**Figure 1.**
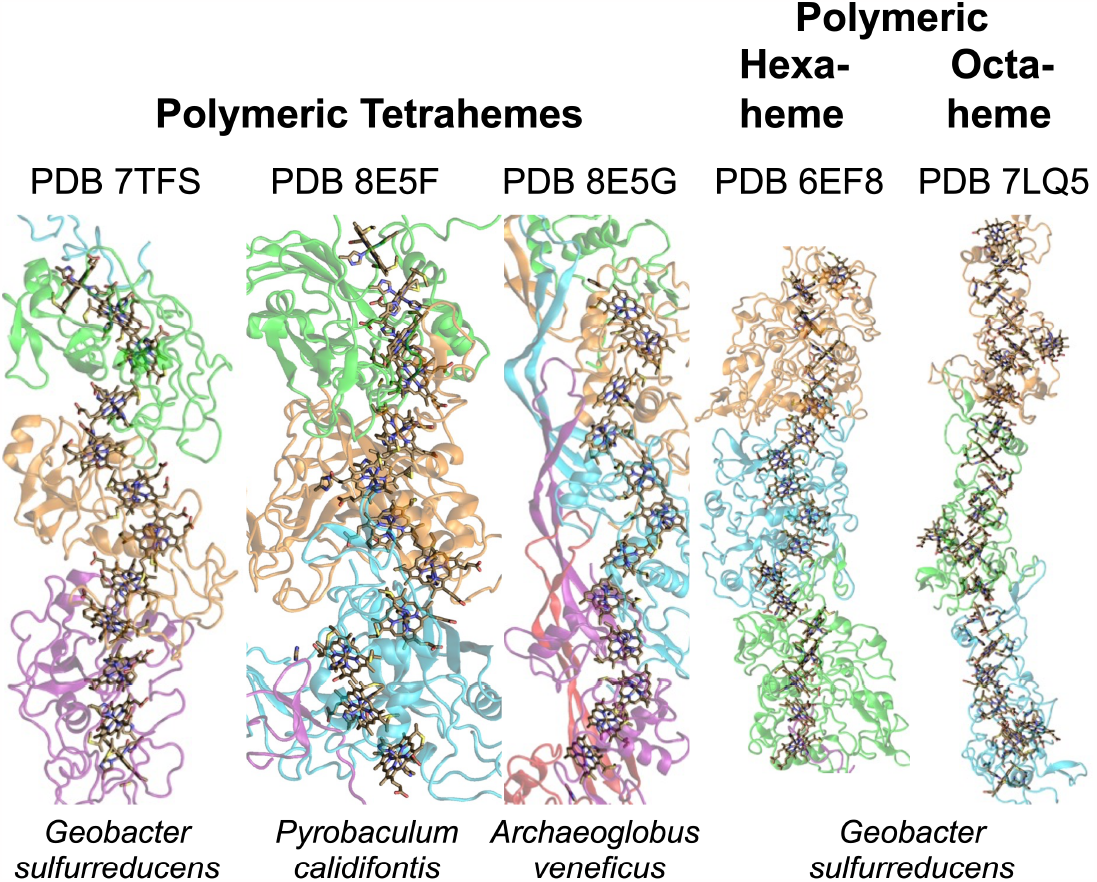
Structural models of the cytochrome ‘nanowires’ assessed for redox conductivity using the python program BioDC in the present work. Cyan, green, orange, lavender, and red indicate chains A through E, respectively, in the indicated PDBs. Note that PDBs 6NEF and 8D9M, which are of the same proteins reported in PDBs 6EF8 and 7LQ5, respectively, were also included in the computations for comparison. All models were typically kept to trimeric assembly, but in every case except PDB 7LQ5 (and 8D9M), a portion of a fourth and/or fifth subunit needed to be included to provide inter-domain axial His ligands to the hemes within the trimer unit.

Redox conduction in cytochrome filaments is pictured as a cascade of reduction-oxidation (redox) reactions between adjacent hemes, which may be facilitated in some cases by covalent protein backbone or inter-heme H-bonds.^15^ In the redox conduction mechanism, a mobile electron resides on a heme until a rare thermal fluctuation brings the heme into energetic degeneracy with a neighboring heme. At that instant, the electron may (but not always) tunnel (or ‘hop’) to the neighboring heme, where it waits for another rare thermal fluctuation to repeat the process so it can proceed along the chain. The rate of each heme-to-heme electron transfer (k_et_) is expected to be well-described by standard non-adiabatic Marcus theory (Eq. 1),^21, 22^ where ⟨H⟩, λ, and ΔG^∘^ are, respectively, the donor-acceptor electronic coupling, the reorganization energy, and the reaction free energy. The set of k_et_s define a charge diffusion coefficient (D),^23^ which in turn can be used to simulate Ohmic redox current (Eq. 2).^14, 24^

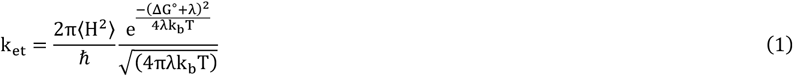

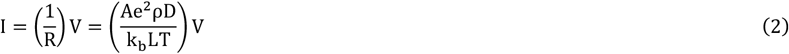

In Eq. 2, R and V are the resistance and voltage of the conductor; A and L are the cross-sectional area and length of the conduction channel; e, k_b_ and T are the elementary charge, the Boltzmann constant, and absolute temperature; and ρ is the charge density. In the present work, the analytical expression derived by Derrida for diffusion along a one-dimensional chain of hopping sites^23^ is used to compute D.

The three energetic parameters for each k_et_ can be computed from quantum mechanical (QM), molecular mechanical (MM), or hybrid QM/MM methodologies.^25^ Typically, hundreds of nanoseconds of classical molecular dynamical (MD) simulations and/or thousands of QM energy evaluations on thermally sampled configurations (QM/MM@MD) are needed to obtain converged quantities.^14^ The computational burden grows rapidly with the number of hemes serving as charge hopping sites in the periodic ‘unit cell’ of the filament. The periodic ‘unit cell’ is usually regarded as the subunit of the homopolymer for computational tractability^14, 15, 18, 26^ but may also refer to the collection of subunits that form one helical pitch of the filament. The influence of the helical environment on redox conductivity has not yet been assessed.

The computational burden becomes even more intractable when the goal is to compare several cytochrome filaments, as in the present Letter. To remove this bottleneck, the BioDC Python program is herein reported as an efficient way to compute redox currents in cytochrome ‘nanowires’ according to Eqs. 1 and 2 and applied to the structures in Figure 1.

BioDC is written in a highly modular and interactive fashion with three main divisions: (1) Structure Preparation and Relaxation (SP&R), (2) Energetic Evaluation (EE), and (3) Redox Current Prediction (RCP). These modules can be run independently or sequentially as a workflow (Figure 2). The interactive BioDC session to compute redox current in OmcS (PDB 6EF8) is provided in Section S1 of the Supporting Information as an example of how the program is used; all the files needed to reproduce the results of this Letter are available at the BioDC GitHub repository.

**Figure 2.**
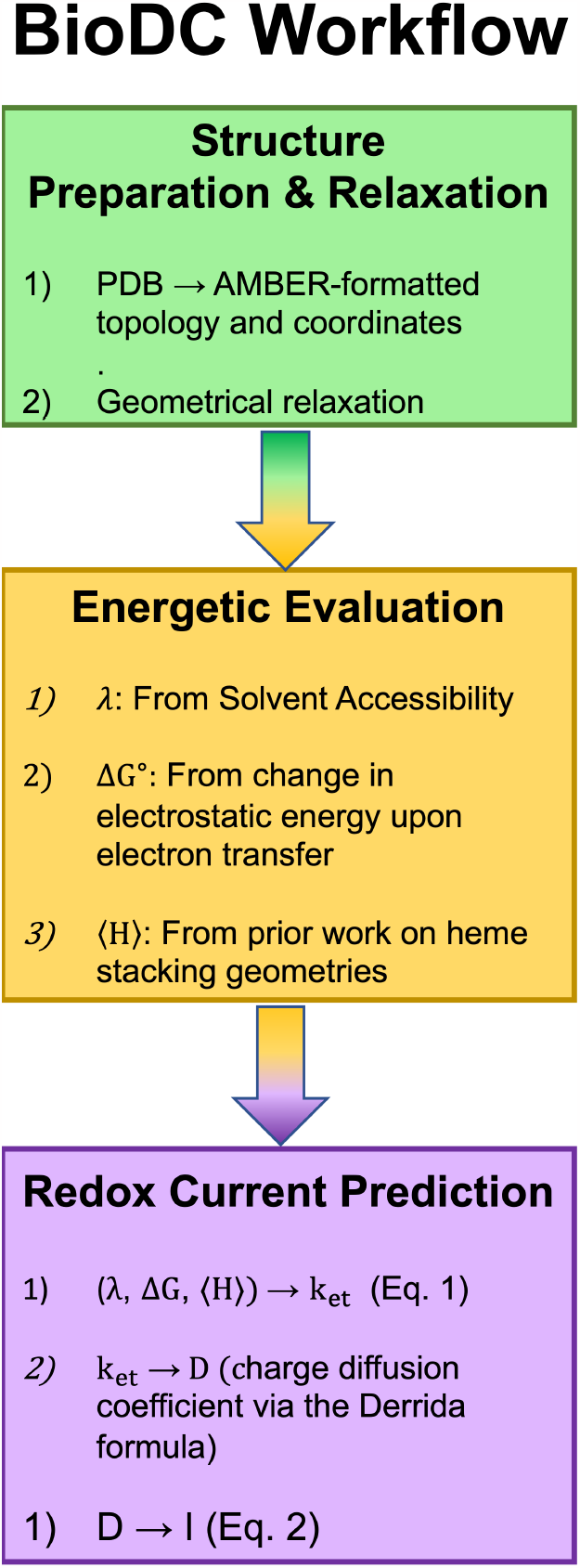
Workflow of the BioDC Python program for computing redox conductivity in polymeric multi-heme cytochromes.

The SP&R module converts a properly prepared structure (see Section S2 of the Supporting Information for details) of a cytochrome ‘nanowire’ in PDB format into Assisted Model Building and Energy Refinement (AMBER)-formatted topology and coordinate files, which are then used to energetically relax the geometry. The structure is described with the ff99SB force-field,^27-29^ which has previously been shown to well-reproduce *relative* differences in electric fields^30^ and to predict heme redox potentials within ∼0.12 V of experiment in Constant Redox Molecular dynamics (CEMD)^31^ simulations.

The EE module assesses λ, ΔG^∘^, and ⟨H⟩ for each electron transfer reaction in the (user-specified) linear sequence of heme groups. λ is estimated from a form of the Marcus continuum expression (Eq. 3) where the static dielectric constant (ϵ_s_) was parameterized by Blumberger and co-workers^20^ to linearly depend (Eq. 4) on the total (donor + acceptor) solvent accessible surface area (SASA). The parameterization was shown to provide λs that agree well with results from classical MD simulations performed with a polarizable force-field. This approach accounts for the outer-sphere (environmental) contribution to λ, which has been shown to be ≥10-times larger than the inner-sphere component from redox-linked changes in internal degrees of freedom of the heme group.^32^

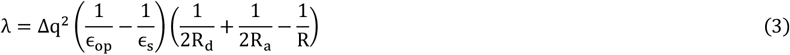

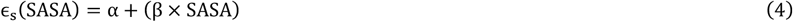

In Eq. 3, the amount of charge transferred is Δq = −1*e*; the optical dielectric constant is taken as ϵ_op_ = 1.84; the radius of the donor or acceptor hemes is R_x_ (x = d or a) = 4.6 Å, and the donor-to-acceptor distance measured between the Fe atoms of the two hemes is typically R_da_= 9 − 12 Å; α and β in Eq. 4 have the values 5.18 and 0.016 Å^-2^.^20^

ΔG^∘^ is estimated from the difference in redox potential between adjacent electron donating 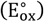 and accepting 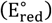 hemes (Eq. 5). In Eq. 5, n is the number of transferred electrons (n = 1) and F is Faraday’s constant (1 eV/V).

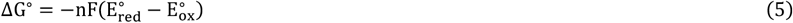

The E^∘^s are estimated from the change in electrostatic energy upon oxidation of each heme in Poisson–Boltzmann Surface Area (PBSA) calculations. This approach was previously used to compute E^∘^s in OmcS from *Geobacter sulfurreducens* and the MtrCAB complex from *Shewanella baltica*.^20^ The neglect of conformational sampling makes the approach considerably more approximate than the use of Multi-Conformational Continuum Electrostatics (MCCE),^33^ CEMD,^31^ or QM/MM@MD^14^ to compute E^∘^. BioDC allows the user to enter energetic parameters computed by these other methods, comes with a helper script to format the relaxed geometry for use with MCCE, and prepares topology and coordinate files that can be directly submitted to CEMD simulations.

⟨H⟩ is assigned to perpendicular (T-stacked) and parallel-displaced (slip-stacked) adjacent heme pairs with values of 2.0 and 8.0 meV, respectively. These values are consistent with those reported previously for heme cofactors in these two highly conserved stacking geometries,^14, 15, 20, 34^ and the finding that environmental effects on ⟨H⟩ can be neglected.^25^ Alternatively, the ⟨H⟩s can be computed with QM or QM/MM methods and entered by the user during the interactive BioDC session.

The RCP module of BioDC uses the estimated or user-entered values for λ, ΔG^∘^, and ⟨H⟩ to compute Marcus-theory charge transfer rates (Eq. 1) and the redox current (Eq. 2) through a cytochrome ‘nanowire.’ Figure 3 shows the results of the BioDC workflow applied to OmcE, OmcS, OmcZ, A3MW92, and F2KMU8 (see Figure 1 for the structures). The BioDC estimates for λ, ΔG^∘^, and ⟨H⟩ used in Eq 1 to obtain these results are given in Sections S3 – S5 of the Supporting Information; the corresponding results previously reported^15^ from a QM/MM@MD protocol are given for reference in Sections S6 – S8.

**Figure 3.**
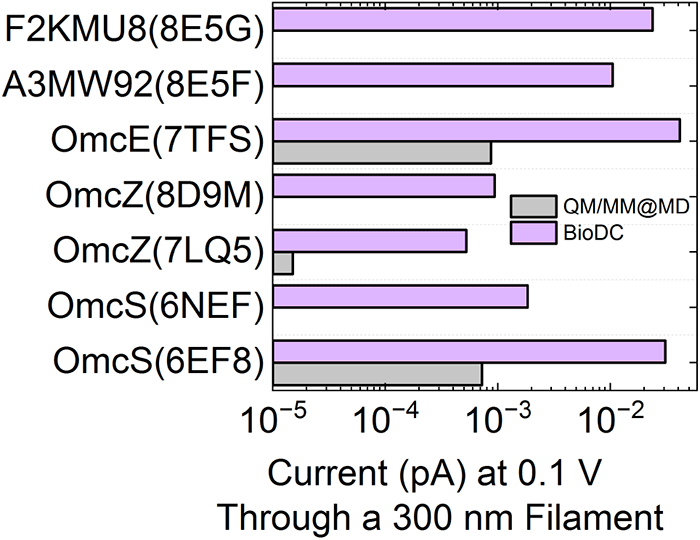
Redox currents in cytochrome ‘nanowires’ predicted by (*grey*) extensive QM/MM energy evaluations at classical MD-generated configurations (QM/MM@MD), and (*lavender*) BioDC. The QM/MM@MD results are reproduced from Ref. 15. The vertical axis gives the name of each protein followed by the Protein Data Bank accession code in parentheses.

A few observations are immediately striking from Figure 3:

1. BioDC predicts redox currents that differ by <10^2^ from those obtained with ∼5000 QM/MM energy evaluations on configurations from 2 μs of classical MD for Omc-E, S, and Z separately.^15^ Figure S1 shows that the deviations in the redox current caused by estimating λ, ΔG^∘^, or ⟨H⟩ with BioDC (and the other two quantities with QM/MM@MD) are typically within a factor of 10 from the full QM/MM@MD result. Given the harsh approximations made in BioDC, and the nearly systematic nature of the discrepancy, BioDC represents a reasonable accuracy-versus-efficiency tradeoff, especially for qualitative or semi-quantitative high-throughput analyses. Interestingly, the redox currents predicted by the more sophisticated QM/MM@MD approach are all *smaller* than those predicted by BioDC.
2. The predicted current through OmcS is 17-times larger for the 6EF8 versus 6NEF cryogenic electron microscopy (CryoEM) structural model, which were both resolved to 3.4 – 3.7 Å. By contrast, the predicted current through OmcZ is only 2-times larger for the 8D9M than 7LQ5 CryoEM model, even though these structures were refined to markedly different (4.2 and 3.4 Å, respectively) resolutions. This up-to 20-fold dependence on the initial CryoEM structural model is useful for putting into context the ∼50-fold difference between BioDC and QM/MM@MD results.
3. Redox currents are similar (within a factor of 4) for polymerized tetra-heme filaments from a heterotrophic, mesophilic proteobacterium (*Geobacter sulfurreducens*, OmcE), a heterotrophic, hyperthermophilic archaeon (*Pyrobaculum calidifontis*, 8E5F), and a chemolithoautotrophic, hyperthermophilic archaeon (*Archaeoglobus veneficus*, 8E5G). Overall, redox currents across all examined filaments vary by a factor of 80, which is only ∼3-times larger than the variation that can arise from the considered CryoEM model.

Thus, there may be, at most, a <10^2^-fold variation in current among all the examined filaments, with OmcZ consistently found to be the least conductive. Note that the computed conductivity for OmcZ is likely an overestimate because no electrons are assumed to be diverted away from the main chain onto the branched, solvent-exposed heme. This result is in stark disagreement with the reported 10^3^-fold greater conductivity of Omc-Z versus S in CP-AFM experiments,^12^ which may be attributable to better heme-electrode contacts because of a fully solvent-exposed heme in OmcZ,^10^ a physiologically irrelevant charge transport mechanism^13^ or an artifact of experimental conditions.^2^

In summary, an efficient computational workflow named BioDC has been implemented, validated, and used to automate and accelerate the computation of redox currents in polymeric multi-heme cytochrome ‘nanowires.’ BioDC delivers results in reasonable and reliable agreement with more sophisticated MD or QM/MM protocols at substantially reduced effort. Leveraging this efficiency for a comparative analysis, BioDC predicts that cytochrome ‘nanowires’ are functionally robust (Figure 3) to structural heterogeneity (Figure 1), suggesting that the protein scaffold can evolve independently of the enclosed heme chain to form a habitat-customized interface for extracellular (microbial or mineral) electron acceptors. The scheme is analogous to the habitat-customized interface provided by light harvesting antennas for the highly conserved photosystems of photosynthesis. Thus, this Letter demonstrates that a meta-principle of evolutionary design applies to cytochrome ‘nanowires’; namely, where possible, re-package, instead of re-invent old solutions to life’s ever-changing challenges.

## Computational Methods

The BioDC program developed and used in the present work is available at the GitHub repository: https://github.com/Mag14011/BioDC. The implemented methodologies are a combination of those reported in Refs. 14 and 20. BioDC calls upon Visual Molecular Dynamics (VMD)^35^ as well as the TLEaP, ParmEd, CPPTRAJ, PMEMD, and PBSA programs of the AMBER Molecular Dynamics suite.^36, 37^ VMD is used to edit the provided PDB and to measure heme solvent accessibility. TLEaP is used to build the AMBER-formatted topology and coordinate files for the inputted structure. ParmEd is used to switch between the force-field definitions for the oxidized or reduced state for each heme. CPPTRAJ is used to ensure that atom/residue numbering for a multi-chain structure conforms to AMBER conventions (*i*.*e*., bonding order), and to measure the angle between the mean macrocycle planes of adjacent hemes. PMEMD is used to energetically relax the structure. PBSA is used to perform Poisson– Boltzmann Surface Area calculations. BioDC also uses a C-program implementation of the Derrida diffusion equation kindly provided by Dr. Fredrik Jansson,^38^ and only modified to read/write files to interface within the BioDC workflow.

## ASSOCIATED CONTENT

### Supporting Information

The following files are available free of charge.

Example interactive session with BioDC to compute redox conductivity; instructions on how to prepare PDBs for use with BioDC; energetic parameters for electron transfer estimated with BioDC and a QM/MM@MD methodology (PDF)

## AUTHOR INFORMATION

### Author Contributions

All original computer code and results reported in this Letter were generated by the corresponding author between 05/23/2023 and 06/06/2023.

### Notes

The author declares no competing financial interest.

## ACKNOWLEDGMENT

Much gratitude is extended to Dr. Fredrik Jansson for permitting the inclusion of his C-program implementation of the Derrida diffusion equation into the BioDC workflow reported here.

## ABBREVIATIONS

AMBER: Assisted Model Building and Energy Refinement
CEMD: Constant Redox Molecular Dynamics
CP-AFM: conducting-probe atomic force microscopy
CryoEM: cryogenic electron microscopy
EE: Energetic Evaluation
MCCE: Multi-Conformational Continuum Electrostatics
PBSA: Poisson-Boltzmann Surface Area
RCP: Redox Current Prediction
SP&R: Structure Preparation and Relaxation
QM: quantum mechanics
/MM: molecular mechanics
QM/MM@MD: quantum mechanical/molecular mechanical computations at MD-generated configurations

## References

(1) Lovley, D. R.; Holmes, D. E. Electromicrobiology: the ecophysiology of phylogenetically diverse electroactive microorganisms. Nat. Rev. Microbiol. 2022, 20 (1), 5–19.

(2) Baquero, D. P.; Cvirkaite-Krupovic, V.; Hu, S. S.; Fields, J. L.; Liu, X.; Rensing, C.; Egelman, E. H.; Krupovic, M.; Wang, F. Extracellular cytochrome nanowires appear to be ubiquitous in prokaryotes. Cell, 2023; Vol. 186, pp 1–12.

(3) Lovley, D. R. Syntrophy goes electric: direct interspecies electron transfer. Annu. Rev. Microbiol. 2017, 71, 643–664.

(4) Jelen, B. I.; Giovannelli, D.; Falkowski, P. G. The role of microbial electron transfer in the coevolution of the biosphere and geosphere. Annu. Rev. Microbiol. 2016, 70, 45–62.

(5) Jiang, Y.; Shi, M.; Shi, L. Molecular underpinnings for microbial extracellular electron transfer during biogeochemical cycling of earth elements. Sci. China: Life Sci. 2019, 62 (10), 1275–1286.

(6) Lovley, D. R.; Yao, J. Intrinsically conductive microbial nanowires for ‘green’electronics with novel functions. Trends Biotechnol. 2021, 39 (9), 940–952.

(7) Wang, F.; Gu, Y.; O’Brien, J. P.; Sophia, M. Y.; Yalcin, S. E.; Srikanth, V.; Shen, C.; Vu, D.; Ing, N. L.; Hochbaum, A. I. Structure of microbial nanowires reveals stacked hemes that transport electrons over micrometers. Cell 2019, 177 (2), 361–369. e310.

(8) Filman, D. J.; Marino, S. F.; Ward, J. E.; Yang, L.; Mester, Z.; Bullitt, E.; Lovley, D. R.; Strauss, M. Cryo-EM reveals the structural basis of long-range electron transport in a cytochrome-based bacterial nanowire. Commun. Biol. 2019, 2 (1), 219.

(9) Wang, F.; Mustafa, K.; Suciu, V.; Joshi, K.; Chan, C. H.; Choi, S.; Su, Z.; Si, D.; Hochbaum, A. I.; Egelman, E. H. Cryo-EM structure of an extracellular Geobacter OmcE cytochrome filament reveals tetrahaem packing. Nat. Microbiol. 2022, 7 (8), 1291–1300.

(10) Wang, F.; Chan, C. H.; Suciu, V.; Mustafa, K.; Ammend, M.; Si, D.; Hochbaum, A. I.; Egelman, E. H.; Bond, D. R. Structure of Geobacter OmcZ filaments suggests extracellular cytochrome polymers evolved independently multiple times. Elife 2022, 11, e81551.

(11) Gu, Y.; Guberman-Pfeffer, M. J.; Srikanth, V.; Shen, C.; Giska, F.; Gupta, K.; Londer, Y.; Samatey, F. A.; Batista, V. S.; Malvankar, N. S. Structure of Geobacter cytochrome OmcZ identifies mechanism of nanowire assembly and conductivity. Nat. Microbiol. 2023, 8 (2), 284–298.

(12) Yalcin, S. E.; O’Brien, J. P.; Gu, Y.; Reiss, K.; Yi, S. M.; Jain, R.; Srikanth, V.; Dahl, P. J.; Huynh, W.; Vu, D. Electric field stimulates production of highly conductive microbial OmcZ nanowires. Nat. Chem. Biol. 2020, 16 (10), 1136–1142.

(13) Futera, Z.; Wu, X.; Blumberger, J. Tunneling-to-Hopping Transition in Multiheme Cytochrome Bioelectronic Junctions. J. Phys. chem. Lett. 2023, 14, 445–452.

(14) Guberman-Pfeffer, M. J. Assessing Thermal Response of Redox Conduction for Anti-Arrhenius Kinetics in a Microbial Cytochrome Nanowire. J. Phys. Chem. B 2022, 126 (48), 10083–10097.

(15) Guberman-Pfeffer, M. J. Structural Determinants of Redox Conduction Favor Robustness over Tunability in Microbial Cytochrome Nanowires. BioRxiv DOI: https://doi.org/10.1101/2023.01.21.525004

(16) Eshel, Y.; Peskin, U.; Amdursky, N. Coherence-assisted electron diffusion across the multi-heme protein-based bacterial nanowire. Nanotechnology 2020, 31 (31), 314002.

(17) Naaman, R.; Waldeck, D. H.; Fransson, J. New Perspective on Electron Transfer through Molecules. J. Phys. Chem. Lett. 2022, 13, 11753–11759.

(18) Livernois, W.; Anantram, M. A Spin-Dependent Model for Multi-Heme Bacterial Nanowires. ACS Nano 2023, 17, 10, 9059–9068

(19) Amdursky, N.; Ferber, D.; Pecht, I.; Sheves, M.; Cahen, D. Redox activity distinguishes solid-state electron transport from solution-based electron transfer in a natural and artificial protein: Cytochrome C and hemin-doped human serum albumin. Phys. Chem. Chem. Phys. 2013, 15 (40), 17142–17149.

(20) Jiang, X.; van Wonderen, J. H.; Butt, J. N.; Edwards, M. J.; Clarke, T. A.; Blumberger, J. Which multi-heme protein complex transfers electrons more efficiently? Comparing MtrCAB from Shewanella with OmcS from Geobacter. J. Phys. Chem. Lett. 2020, 11 (21), 9421–9425.

(21) Marcus, R. A. On the theory of oxidation-reduction reactions involving electron transfer. I. J. Chem. Phys. 1956, 24 (5), 966–978.

(22) Marcus, R. Discussion comment on mixed reaction-diffusion controlled rates. Discuss. Faraday Soc. 1960, 29, 21–31.

(23) Derrida, B. Velocity and diffusion constant of a periodic one-dimensional hopping model. J. Stat. Phys. 1983, 31, 433–450.

(24) Polizzi, N. F.; Skourtis, S. S.; Beratan, D. N. Physical constraints on charge transport through bacterial nanowires. Faraday Discuss. 2012, 155, 43–61.

(25) Blumberger, J. Recent advances in the theory and molecular simulation of biological electron transfer reactions. Chem. Rev. 2015, 115 (20), 11191–11238.

(26) Dahl, P. J.; Yi, S. M.; Gu, Y.; Acharya, A.; Shipps, C.; Neu, J.; O’Brien, J. P.; Morzan, U. N.; Chaudhuri, S.; Guberman-Pfeffer, M. J. A 300-fold conductivity increase in microbial cytochrome nanowires due to temperature-induced restructuring of hydrogen bonding networks. Sci. Adv. 2022, 8 (19), eabm7193.

(27) Hornak, V.; Abel, R.; Okur, A.; Strockbine, B.; Roitberg, A.; Simmerling, C. Comparison of multiple Amber force fields and development of improved protein backbone parameters. Proteins: Struct., Funct., Bioinf. 2006, 65 (3), 712–725.

(28) Crespo, A.; Martí, M. A.; Kalko, S. G.; Morreale, A.; Orozco, M.; Gelpi, J. L.; Luque, F. J.; Estrin, D. A. Theoretical study of the truncated hemoglobin HbN: exploring the molecular basis of the NO detoxification mechanism. J. Am. Chem. Soc. 2005, 127 (12), 4433–4444.

(29) Henriques, J. o.; Costa, P. J.; Calhorda, M. J.; Machuqueiro, M. Charge Parametrization of the DvH-c3 Heme Group: Validation Using Constant-(pH, E) Molecular Dynamics Simulations. J. Phys. Chem. B 2013, 117 (1), 70–82.

(30) Wang, X.; He, X. An Ab Initio QM/MM Study of the Electrostatic Contribution to Catalysis in the Active Site of Ketosteroid Isomerase. Molecules 2018, 23 (10), 2410.

(31) Cruzeiro, V. W. D.; Feliciano, G. T.; Roitberg, A. E. Exploring coupled redox and pH processes with a force-field-based approach: applications to five different systems. J. Am. Chem. Soc. 2020, 142 (8), 3823–3835.

(32) Breuer, M.; Zarzycki, P.; Shi, L.; Clarke, T. A.; Edwards, M. J.; Butt, J. N.; Richardson, D. J.; Fredrickson, J. K.; Zachara, J. M.; Blumberger, J. Molecular structure and free energy landscape for electron transport in the decahaem cytochrome MtrF. Biochem. Soc. Trans. 2012 40, 6, 1198–1203.

(33) Zheng, Z.; Gunner, M. Analysis of the electrochemistry of hemes with Ems spanning 800 mV. Proteins: Struct., Funct., Bioinf. 2009, 75 (3), 719–734.

(34) Jiang, X.; Burger, B.; Gajdos, F.; Bortolotti, C.; Futera, Z.; Breuer, M.; Blumberger, J. Kinetics of trifurcated electron flow in the decaheme bacterial proteins MtrC and MtrF. Proc. Natl. Acad. Sci. U. S. A. 2019, 116 (9), 3425–3430.

(35) Humphrey, W.; Dalke, A.; Schulten, K. VMD: visual molecular dynamics. J. Mol. Graphics 1996, 14 (1), 33–38.

(36) Case, D. A.; Cheatham III, T. E.; Darden, T.; Gohlke, H.; Luo, R.; Merz Jr, K. M.; Onufriev, A.; Simmerling, C.; Wang, B.; Woods, R. J. The Amber biomolecular simulation programs. J. Comput. Chem. 2005, 26 (16), 1668–1688.

(37) Roe, D. R.; Cheatham III, T. E. PTRAJ and CPPTRAJ: software for processing and analysis of molecular dynamics trajectory data. J. Chem. Theory Comput. 2013, 9 (7), 3084–3095.

(38) Nenashev, A.; Jansson, F.; Baranovskii, S.; Österbacka, R.; Dvurechenskii, A.; Gebhard, F. Effect of electric field on diffusion in disordered materials. II. Two-and three-dimensional hopping transport. Phys. Rev. B 2010, 81 (11), 115204.

