## Supporting Information for "From Hot Water to Dry Dirt: Microbes Use Cytochrome ‘Nanowires’ of Similar Conductivity but Different Structure"

**S2. Representative output for an interactive session with BioDC to analyze redox conduction in OmcS (PDB 6EF8)**

The below output from the terminal session has not been edited other than to highlighted user responses in **bold green**, and additional notes in **bold red** front. Note that "rlwrap: Command not found" is a begin message from VMD on the Linux workstation where the session was ran.

```
==================================================================
                    Weclome to BioDC
          A program that automates and accelerates
            the computation of redox currents in
             (polymeric) multi-heme cytochormes

           Written by Matthew J. Guberman-Pfeffer
               Last Updated: 05/28/2023

This research was supported by the National Institute of General
Medical Sciences of the National Institutes of Health under award
1F32GM142247-01A1.
==================================================================

BioDC presents a highly modular workflow that has three
large divisions:
   (1) Structure Preparaiton & Relaxation
   (2) Energetic Estimation
   (3) Redox Current Prediction

Which of these divisions would you like to perform?
(Enter zero "0" to be guided through the entire
workflow.) (0/1/2/3) 0

===================================================================
First: Structure Preparation & Relaxation
===================================================================

This program requires VMD and the AmberTools package to be
in your system's PATH variable. Are they? (yes/no)? y

Good! Now, here are the PDBs in the present direcotry:

6EF8.pdb
6EF8_C.pdb
6EF8_G.pdb
6EF8_preped.pdb
6EF8_trimer_reord.pdb
6EF8_E.pdb
```

```
HEC.pdb
6EF8_trimer.pdb
6EF8_D.pdb
6EF8_B.pdb
prot.pdb
6EF8_F.pdb
6EF8_A.pdb
Prepare6EF8.pdb

 Which PDB would you like to setup
 (omit the .pdb file extension)? 6EF8_preped
 That PDB was found!

 We need to create a file (ResIndexing.txt) that identifies
 the IDs of the Cys and His residues bound to the heme macrocycle.
 Would you like to create it automatically or manually (auto/man)?
auto

 The automated creation of ResIndexing.txt has two parts:
   1) Writting CreateResIndexing.tcl
   2) Submitting the TCL script to Visual Molecular Dynamics (VMD)

 The TCL script identifies the Cys and His residues within a
 distance cutoff of each heme group and assumes that the residues
 found within that cutoff are bonded to that heme.

 What distance threshold would you liek to use?
 (recommended = 2.5 angstroms)                    2.5
[Note, 2.5 Å was used for all PDBs examined in this work except 6NEF,
for which 3.3 Å was used.]

 Please make sure that the correct residues are identified by,
 for example, creating representations in VMD with the residue IDs
 given on each line of the ResIndexing.txt file.

 If the wrong residues are identified, the setup later with TLEaP
 will fail because the bond definitions will be wrong. In this case,
 please correct the residue IDs and save the changes to
 CorrectedResIndexing.txt. When you re-run this python scirpt, the
 CorrectedResIndexing.txt file will be detected and used to replace
 ResIndexing.txt.

rlwrap: Command not found.

 Your ResIndexing file will now be used with VMD
 to create the HEH, HIO, and CYO residues.

 We will write and submit a script called SetupStructure.tcl to
 perform this magic.

 CAUTION: The magic of the script is only as good as the information
 in ResIndexing.txt. If the wrong residue IDs are specified,
```

```
 everything from hereon out will be, put politely, junk!

 Shall we venture forward with SetupStructure.tcl (yes/no)? y
rlwrap: Command not found.

 VMD finished. Please check SetupStructure.log for any erros. You may
 also want to inspect the generated PDBs for the protein,each heme,
 and each heme propionic acid group.

 Now, we need to stitch the edited PDBs of the protein, hemes,
 and heme propionic acid groups into a single PDB. Then, this
 re-constructed PDB of the multi-heme protein will be processed
 with TLEaP of the AmberTools package to generate topology and
 coordinate files.

 Prefix for output parm/rst7 6EF8
 Should the structure be prepared with
 an explicit or implicit solvent (explicit/implicit)? e
  Using a rectangular or an octahedral box (rec/octahed)? rec
  With how much of a solvent buffer (in angstroms)? 15
  And how many Na+ ions; 0 = enough for charge neutrality? 0
  And how many Cl- ions; 0 = enough for charge neutrality? 0

 The re-compiled structure will now be processed with TLEaP.

 TLEaP finished!
 Please inspect the structure to make sure it is correct.

 Found 6EF8.prmtop and 6EF8.rst7
 Preparing to relax the geometry
 Running minimization ...
 Minimization finished!

 ===================================================================
 Second: Energetic Estimation
 ===================================================================


 Is your structure polymeric (yes/no)? y

 The structure preparation stage required you to place all the heme
 residues from all the chains at the end of the PDB with sequential
 numbering. The programs in AmberTools that will be used to estimate
 the charge transfer energetics want instead the residues to come in
 the order of the connectiviity; that is, the hemes of chain A should
 come both any residue in chain B.

 To oblige this different numbering convention, we'll use the
 CPPTRAJ of the AmberTools package to re-order the residues.
 This process will write a new topology and coordinate file,
 where the latter is of the structure you previously minimized.
```

```
To compute the energetics for heme-to-heme electron transfer. We
need to know the linear sequence of hemes that will serve as
charge hopping sites. Typically the linear sequence is NOT the
sequence of residue IDs in the PDB. We therefore need to specify
the linear sequence to compute the right electron transfer steps.

Linear Sequence: 1275 1260 1263 1269 1266 1272 1293
-------------------------------------------------------------------

Should we compute the reorganization energy (yes/no)? y
Found 6EF8_reord.prmtop and min.rst7

To estimate the reorganization energy (lambda) from the
solvent accessible surface area, two steps will be take:
   (1) Convert min.rst7 to a PDB-formatted file using ambpdb
       Note: This step is skipped because you indicated you
       have a polymeric structute, When the topology was
       re-ordered to conform to AMBER conventions for
       multi-chain structures, min.pdb was already
       created.

   (2) Write and submit a TCL script to VMD

 Now using VMD to compute SASA Donor = 1275 & Acceptor = 1260...
rlwrap: Command not found.
 Now using VMD to compute SASA Donor = 1260 & Acceptor = 1263...
rlwrap: Command not found.
 Now using VMD to compute SASA Donor = 1263 & Acceptor = 1269...
rlwrap: Command not found.
 Now using VMD to compute SASA Donor = 1269 & Acceptor = 1266...
rlwrap: Command not found.
 Now using VMD to compute SASA Donor = 1266 & Acceptor = 1272...
rlwrap: Command not found.
 Now using VMD to compute SASA Donor = 1272 & Acceptor = 1293...
rlwrap: Command not found.
 Computing Reorganization Energy from Solvent Accessibility...
 Done!
-------------------------------------------------------------------

Should we compute the reaction free energy (yes/no)? y

Two different methods are implemented to estimate heme redox
potentials and thereby reaction free energies when an explict
solvent is presnet:

   (1) Compute the change in electrostatic interaciton energy upon
       heme oxidaiton using the Linear Interaction Energy method
       in CPPTRAJ.

   (2) Compute the change in electrostatic interaciton energy upon
       heme oxidaiton using the Poisson-Boltzmann Surface Area
       method implemented in AmberTools (essentially a Delphi-type
```

```
        calculation). In this case, the explicit solvent
        prepared with the sturcutre is discarded.

    Note that method #1 has two advantages:
        (1) It is considerably faster than method #2

        (2) The overall change in electorstatic energy is decomposed
            into contributions from different groups of residues.

 Should we use the LIE or PBSA method (lie/pbsa)? pbsa

 Generating topology for oHEH-1275 ...

 Generating topology for rHEH-1275 ...

 Generating topology for oHEH-1260 ...

 Generating topology for rHEH-1260 ...

 Generating topology for oHEH-1263 ...

 Generating topology for rHEH-1263 ...

 Generating topology for oHEH-1269 ...

 Generating topology for rHEH-1269 ...

 Generating topology for oHEH-1266 ...

 Generating topology for rHEH-1266 ...

 Generating topology for oHEH-1272 ...

 Generating topology for rHEH-1272 ...

 Generating topology for oHEH-1293 ...

 Generating topology for rHEH-1293 ...

 Submitting 14 PBSA calculations in parallel
  Finished: <Popen: returncode: 0 args: 'pbsa -O -i pbsa.key -o
pbsa_o1275 -p o1275.prmt...>
  Finished: <Popen: returncode: 0 args: 'pbsa -O -i pbsa.key -o
pbsa_r1275 -p r1275.prmt...>
  Finished: <Popen: returncode: 0 args: 'pbsa -O -i pbsa.key -o
pbsa_o1260 -p o1260.prmt...>
  Finished: <Popen: returncode: 0 args: 'pbsa -O -i pbsa.key -o
pbsa_r1260 -p r1260.prmt...>
  Finished: <Popen: returncode: 0 args: 'pbsa -O -i pbsa.key -o
pbsa_o1263 -p o1263.prmt...>
  Finished: <Popen: returncode: 0 args: 'pbsa -O -i pbsa.key -o
pbsa_r1263 -p r1263.prmt...>
```

```
  Finished: <Popen: returncode: 0 args: 'pbsa -O -i pbsa.key -o
pbsa_o1269 -p o1269.prmt...>
  Finished: <Popen: returncode: 0 args: 'pbsa -O -i pbsa.key -o
pbsa_r1269 -p r1269.prmt...>
  Finished: <Popen: returncode: 0 args: 'pbsa -O -i pbsa.key -o
pbsa_o1266 -p o1266.prmt...>
  Finished: <Popen: returncode: 0 args: 'pbsa -O -i pbsa.key -o
pbsa_r1266 -p r1266.prmt...>
  Finished: <Popen: returncode: 0 args: 'pbsa -O -i pbsa.key -o
pbsa_o1272 -p o1272.prmt...>
  Finished: <Popen: returncode: 0 args: 'pbsa -O -i pbsa.key -o
pbsa_r1272 -p r1272.prmt...>
  Finished: <Popen: returncode: 0 args: 'pbsa -O -i pbsa.key -o
pbsa_o1293 -p o1293.prmt...>
  Finished: <Popen: returncode: 0 args: 'pbsa -O -i pbsa.key -o
pbsa_r1293 -p r1293.prmt...>

 Result:
  step=0 HEH-1275 EtotOx=-1102.399 eV EtotRed=-1102.067 eV DEtot=-
0.331 eV EelecOx=-753.059 eV EelecRed=-750.648 eV DEelec=-2.411 eV
  step=1 HEH-1260 EtotOx=-1102.399 eV EtotRed=-1101.980 eV DEtot=-
0.419 eV EelecOx=-753.059 eV EelecRed=-750.284 eV DEelec=-2.775 eV
  step=2 HEH-1263 EtotOx=-1102.399 eV EtotRed=-1101.996 eV DEtot=-
0.402 eV EelecOx=-753.059 eV EelecRed=-750.264 eV DEelec=-2.795 eV
  step=3 HEH-1269 EtotOx=-1102.399 eV EtotRed=-1102.080 eV DEtot=-
0.319 eV EelecOx=-753.059 eV EelecRed=-750.573 eV DEelec=-2.486 eV
  step=4 HEH-1266 EtotOx=-1102.399 eV EtotRed=-1102.068 eV DEtot=-
0.331 eV EelecOx=-753.059 eV EelecRed=-750.556 eV DEelec=-2.503 eV
  step=5 HEH-1272 EtotOx=-1102.399 eV EtotRed=-1102.051 eV DEtot=-
0.347 eV EelecOx=-753.059 eV EelecRed=-750.643 eV DEelec=-2.416 eV
  step=6 HEH-1293 EtotOx=-1102.399 eV EtotRed=-1102.083 eV DEtot=-
0.316 eV EelecOx=-753.059 eV EelecRed=-750.779 eV DEelec=-2.280 eV
 ------------------------------------------------------------------

 Should we estimate the electronic coupling from the geometry
(yes/no)? y

 Assiging coupling values based on inter-macrocycle planar anlge ...
 Done!
 ------------------------------------------------------------------

 Should we compute the non-adiabatic Marcus-theory rates (yes/no)? y

 Step #0:
    Activation Energy:
        Forward: 3.122E-01 eV
        Reverse: 2.247E-01 eV
    Rates:
        Forward: 3.685E+05
        Reverse: 1.088E+07

 Step #1:
```

```
   Activation Energy:
       Forward: 2.551E-01 eV
       Reverse: 2.718E-01 eV
   Rates:
       Forward: 5.405E+07
       Reverse: 2.831E+07

Step #2:
   Activation Energy:
       Forward: 2.240E-01 eV
       Reverse: 3.074E-01 eV
   Rates:
       Forward: 1.127E+07
       Reverse: 4.461E+05

Step #3:
   Activation Energy:
       Forward: 2.903E-01 eV
       Reverse: 2.783E-01 eV
   Rates:
       Forward: 1.337E+07
       Reverse: 2.126E+07

Step #4:
   Activation Energy:
       Forward: 2.985E-01 eV
       Reverse: 2.821E-01 eV
   Rates:
       Forward: 6.007E+05
       Reverse: 1.133E+06

Step #5:
   Activation Energy:
       Forward: 2.484E-01 eV
       Reverse: 2.795E-01 eV
   Rates:
       Forward: 7.020E+07
       Reverse: 2.106E+07

====================================================================
Third: Redox Current Prediction
====================================================================
```

This division of the BioDC workflow computes, via the analytical Derrida formula, the charge diffuction coefficient based on the non-adiabatic Marcus theory rates. The diffusion coefficient is then related to the electrical resistance and used in Ohm's law to compute the current as a function of applied bias. Note that this approach is only rigorously correct in the limit of zero bias.

Found rates.txt, which is needed to proceed!

```
 We will now compile and run a C-program kindly provided by
 Dr. Fredrik Jansson that has been modified only to
 interface I/O operations with the BioDC program
cc  -Wall  -g  -O3  -ffast-math   -march=native  -fomit-frame-pointer  -
std=c99    derrida.c   -o derrida
The are 6 electron transfer steps
3.685000E+05 2.106419E+07
5.405000E+07 1.087768E+07
1.127000E+07 2.830783E+07
1.337000E+07 4.461479E+05
6.007000E+05 2.126395E+07
7.020000E+07 1.133267E+06
V: -164726.788445
D: 4.184524e-09

 Found D.txt, which is needed to proceed!
 We will at last compute the redox current.
 To do this, some system-specific information is needed.

 Please provide the following parmaeters:
  Temperature (K)? 300
  Number of Charges per subunit? 3
```

**[Note, the number of charges per subunit was 2, 3, or 4, respectively, for tetrahemes, hexahemes, and octahemes, respectively.]**

```
rlwrap: Command not found.

 The length of a subunit of the cytochrome polymer is needed.
  The subunit length measured between the first and the last heme
  specified in LinearizedHemeSequence.txt is 4.91E-07

  Length of wire (cm)? 3E-5
  Experimental Conductance (S/cm) ?
   (Enter "0" if not known)          2.67E-10
```

**[Note, Value comes form *Nanotechnology* 2020, *31* (31), 314002.]**

```
 Charge per Subunit Length            = 6.115804e+06 q/cm
 Cross-Sectional Area                 = 1.767146e-14 cm^2
 Charge Density                       = 3.460837e+20 q/cm^2
 Experimental Diffusion Constant      = 2.112780e-04 cm^2/s
 Computed Diffusion Constant          = 4.184524e-09 cm^2/s

 Voltage (V) Exp. Current (pA) Computed Current (pA)
      -0.500          -133.500             -2.644E-03
      -0.450          -120.150             -2.380E-03
      -0.400          -106.800             -2.115E-03
      -0.350           -93.450             -1.851E-03
      -0.300           -80.100             -1.586E-03
      -0.250           -66.750             -1.322E-03
      -0.200           -53.400             -1.058E-03
      -0.150           -40.050             -7.932E-04
```

```
      -0.100          -26.700          -5.288E-04
      -0.050          -13.350          -2.644E-04
      -0.000           -0.000          -5.871E-19
       0.050           13.350           2.644E-04
       0.100           26.700           5.288E-04
       0.150           40.050           7.932E-04
       0.200           53.400           1.058E-03
       0.250           66.750           1.322E-03
       0.300           80.100           1.586E-03
       0.350           93.450           1.851E-03
       0.400          106.800           2.115E-03
       0.450          120.150           2.380E-03
Done!
```

### S2. How to properly prepare a PDB to be analyzed for redox conduction with BioDC

BioDC has three requirements for the initial PDB: (1) All the protein residues must come before all the heme residues; (2) The residues should be numbered sequentially across all protein chains; and (3) each heme needs to be of the *c*-type, bis-histidine variety.

The third requirement is not strictly necessary. If, for example, you have force-field parameters for a different type of heme, BioDC can still analyze redox conduction so long as you edit the `tleap.in` file written by BioDC to specify the force-field files. But a consideration that must be kept in mind for the cytochrome 'nanowires' is that there may be one or more His ligands coordinated to hemes in the adjacent subunit of the filament. In this case, one or more hemes may be under-coordinated at the termini of the filament, violating the third condition for using BioDC. The solution to this problem is to include a fragment of the subunit donating the coordinating ligand. The fragment should preferably be large enough so that the solvent exposure of the subunit being capped is not too different from if the entire coordinating subunit was included. In the case of OmcS, for example, the first 20 residues of chain D were included to cap the adjacent chain in the subunit to provide the His-16 ligand to the penultimate heme in chain B.

The below BASH script was used to download and edit the structure of OmcS (PDB 6EF8) from the Protein Data Bank so it could be submitted to BioDC. Analogous scripts used for the other structures analyzed in this study are available at the GitHub repository for BioDC (https://github.com/Mag14011/BioDC). Note that the BASH script makes use of the pdb_ttools (available on GitHub (https://github.com/haddocking/pdb-tools) and Visual Molecular Dynamics (VMD). The generated `6EF8_preped.pdb` can be directly submitted to BioDC.

```
#!/bin/bash

pdb_fetch -biounit 6EF8 > 6EF8.pdb

pdb_splitchain 6EF8.pdb
cat 6EF8_D.pdb 6EF8_B.pdb 6EF8_A.pdb 6EF8_C.pdb > 6EF8_trimer.pdb

rm EditPDB.tcl 2> /dev/null

cat >> EditPDB.tcl <<- EOF
mol new 6EF8_trimer.pdb
set prot [atomselect top "(chain A B C and not resname HEC) or (chain
D and resid 1 to 20)"]
set HEC  [atomselect top "chain A B C and resname HEC"]
\$prot writepdb prot.pdb
\$HEC writepdb HEC.pdb
exit
EOF

vmd -e EditPDB.tcl > EditPDB.log

cat prot.pdb HEC.pdb | egrep -v "CRYST1|END" > 6EF8_trimer_reord.pdb
pdb_reres 6EF8_trimer_reord.pdb > 6EF8_trimer_reord_reres.pdb
mv 6EF8_trimer_reord_reres.pdb 6EF8_preped.pdb
```

S3. Reorganization energies estimated from heme solvent accessibility using BioDC

| Heme Pair | Total SASA ($Å^2$) | $\epsilon_s$ | $R_{da}$ (Å) | λ (eV) |
|---|---|---|---|---|
| A3MW92 (PDB 8E5F) | | | | |
| 942 → 939 | 29.6 | 5.65 | 11.4 | 0.69 |
| 939 → 936 | 51.9 | 6.01 | 9.4 | 0.60 |
| 936 → 933 | 41.3 | 5.84 | 12.3 | 0.73 |
| 933 → 954 | 17.8 | 5.47 | 10.3 | 0.62 |
| F2KMU8 (PDB 8E5G) | | | | |
| 1122 → 1125 | 27.8 | 5.62 | 9.4 | 0.59 |
| 1125 → 1128 | 30.0 | 5.66 | 11.9 | 0.70 |
| 1128 → 1131 | 16.4 | 5.44 | 9.6 | 0.59 |
| 1131 → 1134 | 15.1 | 5.42 | 11.9 | 0.69 |
| OmcS (PDB 6EF8) | | | | |
| 1275 → 1260 | 7.29 | 5.30 | 11.3 | 0.660 |
| 1260 → 1263 | 5.81 | 5.27 | 9.2 | 0.553 |
| 1263 → 1269 | 1.03 | 5.20 | 11.4 | 0.656 |
| 1269 → 1266 | 64.9 | 6.22 | 8.8 | 0.572 |
| 1266 → 1272 | 69.4 | 6.30 | 12.5 | 0.761 |
| 1272 → 1293 | 6.6 | 5.29 | 9.1 | 0.550 |
| OmcS (PDB 6NEF) | | | | |
| 1260 → 1263 | 81.1 | 6.47 | 12.5 | 0.77 |
| 1263 → 1266 | 69.5 | 6.29 | 9.2 | 0.60 |
| 1266 → 1269 | 2.8 | 5.23 | 11.4 | 0.66 |
| 1269 → 1272 | 6.4 | 5.28 | 9.1 | 0.54 |
| 1272 → 1275 | 10.1 | 5.34 | 11.4 | 0.67 |
| 1275 → 1242 | 17.0 | 5.45 | 9.4 | 0.57 |
| OmcZ (PDB 7LQ5) | | | | |
| 562 → 553 | 10.2 | 5.34 | 11.3 | 0.66 |
| 553 → 559 | 8.3 | 5.31 | 9.0 | 0.55 |
| 559 → 550 | 5.3 | 5.27 | 9.8 | 0.59 |
| 550 → 541 | 119.0 | 7.08 | 9.7 | 0.66 |
| 541 → 547 | 143.1 | 7.47 | 8.6 | 0.60 |
| 547 → 544 | 32.4 | 5.70 | 9.7 | 0.61 |
| 544 → 844 | 9.3 | 5.33 | 9.2 | 0.56 |
| OmcZ (PDB 8D9M) | | | | |
| 562 → 559 | 24.5 | 5.57 | 11.4 | 0.68 |
| 559 → 553 | 26.8 | 5.60 | 9.2 | 0.57 |
| 553→ 550 | 28.3 | 5.63 | 10.3 | 0.63 |
| 550 → 547 | 138.9 | 7.40 | 9.9 | 0.69 |
| 547 → 544 | 139.9 | 7.42 | 8.9 | 0.62 |
| 544 → 541 | 40.5 | 7.53 | 9.9 | 0.63 |

| | | | | |
|---|---|---|---|---|
| 541 → 844 | 32.2 | 5.70 | 11.4 | 0.56 |

**S4. Redox potentials and reaction free energies estimated with BioDC from changes in electrostatic energy upon oxidation/reduction in Poisson-Boltzmann Surface Area calculations**

<u>A3MW92 (PDB 8E5F):</u>

(HEH-942 = -0.225 eV) -> (HEH-939 = -0.366 eV); ΔG = 0.141 eV
(HEH-939 = -0.366 eV) -> (HEH-936 = -0.257 eV); ΔG = -0.109 eV
(HEH-936 = -0.257 eV) -> (HEH-933 = -0.271 eV); ΔG = 0.014 eV
(HEH-933 = -0.271 eV) -> (HEH-954 = -0.226 eV); ΔG = -0.044 eV

<u>F2KMU8 (PDB 8E5G):</u>

(HEH-1122 = -0.458 eV) -> (HEH-1125 = -0.481 eV); DG = 0.023 eV
(HEH-1125 = -0.481 eV) -> (HEH-1128 = -0.514 eV); DG = 0.033 eV
(HEH-1128 = -0.514 eV) -> (HEH-1131 = -0.586 eV); DG = 0.072 eV
(HEH-1131 = -0.586 eV) -> (HEH-1134 = -0.496 eV); DG = -0.090 eV

<u>OmcE (PDB 7TFS):</u>

(HEH-651 = -0.364 eV) -> (HEH-654 = -0.338 eV); ΔG = -0.027 eV
(HEH-654 = -0.338 eV) -> (HEH-657 = -0.472 eV); ΔG = 0.134 eV
(HEH-657 = -0.472 eV) -> (HEH-660 = -0.458 eV); ΔG = -0.014 eV
(HEH-660 = -0.458 eV) -> (HEH-639 = -0.376 eV); ΔG = -0.082 eV

<u>OmcS (PDB 6EF8):</u>

(HEH-1275 = -0.331 eV) -> (HEH-1260 = -0.419 eV); ΔG = 0.088 eV
(HEH-1260 = -0.419 eV) -> (HEH-1263 = -0.402 eV); ΔG = -0.017 eV
(HEH-1263 = -0.402 eV) -> (HEH-1269 = -0.319 eV); ΔG = -0.083 eV
(HEH-1269 = -0.319 eV) -> (HEH-1266 = -0.331 eV); ΔG = 0.012 eV
(HEH-1266 = -0.331 eV) -> (HEH-1272 = -0.347 eV); ΔG = 0.016 eV
(HEH-1272 = -0.347 eV) -> (HEH-1293 = -0.316 eV); ΔG = -0.031 eV

<u>OmcS (PDB 6NEF):</u>

(HEH-1260 = -0.303 eV) -> (HEH-1263 = -0.153 eV); ΔG = -0.150 eV
(HEH-1263 = -0.153 eV) -> (HEH-1266 = -0.163 eV); ΔG = 0.011 eV
(HEH-1266 = -0.163 eV) -> (HEH-1269 = -0.289 eV); ΔG = 0.126 eV
(HEH-1269 = -0.289 eV) -> (HEH-1272 = -0.325 eV); ΔG = 0.035 eV
(HEH-1272 = -0.325 eV) -> (HEH-1275 = -0.344 eV); ΔG = 0.019 eV
(HEH-1275 = -0.344 eV) -> (HEH-1242 = -0.305 eV); ΔG = -0.039 eV

OmcZ (PDB 7LQ5):

(HEH-562 = -0.442 eV) -> (HEH-553 = -0.718 eV); ΔG = 0.276 eV
(HEH-553 = -0.718 eV) -> (HEH-559 = -0.584 eV); ΔG = -0.134 eV
(HEH-559 = -0.584 eV) -> (HEH-550 = -0.590 eV); ΔG = 0.006 eV
(HEH-550 = -0.590 eV) -> (HEH-541 = -0.404 eV); ΔG = -0.186 eV
(HEH-541 = -0.404 eV) -> (HEH-547 = -0.366 eV); ΔG = -0.038 eV
(HEH-547 = -0.366 eV) -> (HEH-544 = -0.330 eV); ΔG = -0.036 eV
(HEH-544 = -0.330 eV) -> (HEH-844 = -0.457 eV); ΔG = 0.128 eV

OmcZ (PDB 8D9M):

(HEH-562 = -0.474 eV) -> (HEH-559 = -0.678 eV); ΔG = 0.204 eV
(HEH-559 = -0.678 eV) -> (HEH-553 = -0.635 eV); ΔG = -0.043 eV
(HEH-553 = -0.635 eV) -> (HEH-550 = -0.439 eV); ΔG = -0.196 eV
(HEH-550 = -0.439 eV) -> (HEH-547 = -0.441 eV); ΔG = 0.002 eV
(HEH-547 = -0.441 eV) -> (HEH-544 = -0.365 eV); ΔG = -0.077 eV
(HEH-544 = -0.365 eV) -> (HEH-541 = -0.345 eV); ΔG = -0.020 eV
(HEH-541 = -0.345 eV) -> (HEH-844 = -0.464 eV); ΔG = 0.120 eV

**S5. Assignment of electronic couplings from heme stacking geometry using BioDC**

A3MW92 (PDB 8E5F):

Hda(HEH-942 <-> HEH-939) ang. = 109.688 deg.; Hda = 2.000 meV
Hda(HEH-939 <-> HEH-936) ang. = 12.651 deg.; Hda = 8.000 meV
Hda(HEH-936 <-> HEH-933) ang. = 114.051 deg.; Hda = 2.000 meV
Hda(HEH-933 <-> HEH-954) ang. = 5.672 deg.; Hda = 8.000 meV

A3MW92 (PDB 8E5G):

Hda(HEH-1122 <-> HEH-1125) ang. = 15.358 deg.; Hda = 8.000 meV
Hda(HEH-1125 <-> HEH-1128) ang. = 112.297 deg.; Hda = 2.000 meV
Hda(HEH-1128 <-> HEH-1131) ang. = 13.491 deg.; Hda = 8.000 meV
Hda(HEH-1131 <-> HEH-1134) ang. = 116.152 deg.; Hda = 2.000 meV

OmcE (PDB 7TFS):

Hda(HEH-651 <-> HEH-654) ang. = 102.421 deg.; Hda = 2.000 meV
Hda(HEH-654 <-> HEH-657) ang. = 19.754 deg.; Hda = 8.000 meV
Hda(HEH-657 <-> HEH-660) ang. = 81.318 deg.; Hda = 2.000 meV
Hda(HEH-660 <-> HEH-639) ang. = 14.675 deg.; Hda = 8.000 meV

OmcS (PDB 6EF8):

Hda(HEH-1275 <-> HEH-1260) ang. = 94.000 deg.; Hda = 2.000 meV
Hda(HEH-1260 <-> HEH-1263) ang. = 6.063 deg.; Hda = 8.000 meV
Hda(HEH-1263 <-> HEH-1269) ang. = 99.841 deg.; Hda = 2.000 meV
Hda(HEH-1269 <-> HEH-1266) ang. = 9.653 deg.; Hda = 8.000 meV
Hda(HEH-1266 <-> HEH-1272) ang. = 106.091 deg.; Hda = 2.000 meV
Hda(HEH-1272 <-> HEH-1293) ang. = 2.025 deg.; Hda = 8.000 meV

OmcS (PDB 6NEF):

Hda(HEH-1260 <-> HEH-1263) ang. = 104.224 deg.; Hda = 2.000 meV
Hda(HEH-1263 <-> HEH-1266) ang. = 8.741 deg.; Hda = 8.000 meV
Hda(HEH-1266 <-> HEH-1269) ang. = 103.733 deg.; Hda = 2.000 meV
Hda(HEH-1269 <-> HEH-1272) ang. = 12.553 deg.; Hda = 8.000 meV
Hda(HEH-1272 <-> HEH-1275) ang. = 92.629 deg.; Hda = 2.000 meV
Hda(HEH-1275 <-> HEH-1242) ang. = 4.813 deg.; Hda = 8.000 meV

OmcZ (PDB 7LQ5):

Hda(HEH-562 <-> HEH-553) ang. = 105.461 deg.; Hda = 2.000 meV
Hda(HEH-553 <-> HEH-559) ang. = 12.892 deg.; Hda = 8.000 meV
Hda(HEH-559 <-> HEH-550) ang. = 19.806 deg.; Hda = 8.000 meV
Hda(HEH-550 <-> HEH-541) ang. = 96.686 deg.; Hda = 2.000 meV
Hda(HEH-541 <-> HEH-547) ang. = 11.127 deg.; Hda = 8.000 meV
Hda(HEH-547 <-> HEH-544) ang. = 115.305 deg.; Hda = 2.000 meV
Hda(HEH-544 <-> HEH-844) ang. = 20.933 deg.; Hda = 8.000 meV

OmcZ (PDB 8D9M):

| | | | |
|---|---|---|---|
| Hda(HEH-562 <-> HEH-559) ang. = | 102.634 deg.; | Hda = | 2.000 meV |
| Hda(HEH-559 <-> HEH-553) ang. = | 8.823 deg.; | Hda = | 8.000 meV |
| Hda(HEH-553 <-> HEH-550) ang. = | 26.936 deg.; | Hda = | 8.000 meV |
| Hda(HEH-550 <-> HEH-547) ang. = | 104.264 deg.; | Hda = | 2.000 meV |
| Hda(HEH-547 <-> HEH-544) ang. = | 11.072 deg.; | Hda = | 8.000 meV |
| Hda(HEH-544 <-> HEH-541) ang. = | 118.832 deg.; | Hda = | 2.000 meV |
| Hda(HEH-541 <-> HEH-844) ang. = | 22.662 deg.; | Hda = | 8.000 meV |

**S6. Marcus-theory electron transfer energetic parameters for OmcE. Data reproduced form Ref. 15 in the main text.**

| Heme Pair | $\Delta G^{\circ}$ | $\lambda_{st}$ | $\lambda_{var,f}$ | $\lambda_{var,r}$ | $\lambda_{rxn}$ | $\langle H \rangle$ |
|---|---|---|---|---|---|---|
| 4’-1 | 0.188 | 0.710 | 0.724 | 0.741 | 0.688 | 7.797 |
| 1-2 | -0.038 | 0.931 | 1.044 | 0.928 | 0.879 | 1.472 |
| 2-3 | -0.009 | 0.587 | 0.764 | 0.666 | 0.481 | 13.001 |
| 3-4 | -0.099 | 0.766 | 0.747 | 0.756 | 0.780 | 1.775 |
| 4-1’’ | 0.067 | 0.758 | 0.667 | 0.808 | 0.778 | 4.411 |

**S7. Marcus-theory electron transfer energetic parameters for OmcS. Data reproduced form Ref. 15 in the main text.**

| Heme Pair | $\Delta G°$ | $\lambda_{st}$ | $\lambda_{var,f}$ | $\lambda_{var,r}$ | $\lambda_{rxn}$ | $\langle H \rangle$ |
|---|---|---|---|---|---|---|
| 6'-1 | 0.031 | 0.620 | 0.547 | 0.602 | 0.668 | 6.719 |
| 1-2 | 0.014 | 0.946 | 0.869 | 0.952 | 0.983 | 1.131 |
| 2-3 | -0.017 | 0.453 | 0.491 | 0.467 | 0.429 | 6.091 |
| 3-4 | 0.161 | 0.597 | 0.632 | 0.505 | 0.626 | 2.639 |
| 4-5 | -0.063 | 0.659 | 0.635 | 0.499 | 0.765 | 9.520 |
| 5-6 | -0.072 | 0.634 | 0.656 | 0.620 | 0.630 | 1.036 |
| 6-1'' | -0.015 | 0.676 | 0.625 | 0.639 | 0.724 | 7.895 |

**S8. Marcus-theory electron transfer energetic parameters for OmcZ. Data reproduced form Ref. 15 in the main text.**

| Heme Pair | $\Delta G°$ | $\lambda_{st}$ | $\lambda_{var,f}$ | $\lambda_{var,r}$ | $\lambda_{rxn}$ | $\langle H \rangle$ |
|---|---|---|---|---|---|---|
| 7’-1 | 0.189 | 0.671 | 0.826 | 0.693 | 0.593 | 9.826 |
| 1-2 | 0.101 | 0.714 | 0.902 | 0.933 | 0.556 | 2.035 |
| 2-3 | 0.019 | 0.766 | 0.735 | 0.760 | 0.785 | 3.187 |
| 3-4 | 0.163 | 0.869 | 0.894 | 0.798 | 0.892 | 7.150 |
| 4-5 | -0.044 | 0.889 | 1.097 | 0.928 | 0.781 | 4.055 |
| 5-6 | -0.046 | 0.815 | 0.864 | 0.886 | 0.758 | 5.206 |
| 6-7 | -0.258 | 0.777 | 0.767 | 1.010 | 0.680 | 2.323 |
| 7-1’’ | 0.092 | 0.717 | 0.746 | 0.905 | 0.624 | 4.711 |

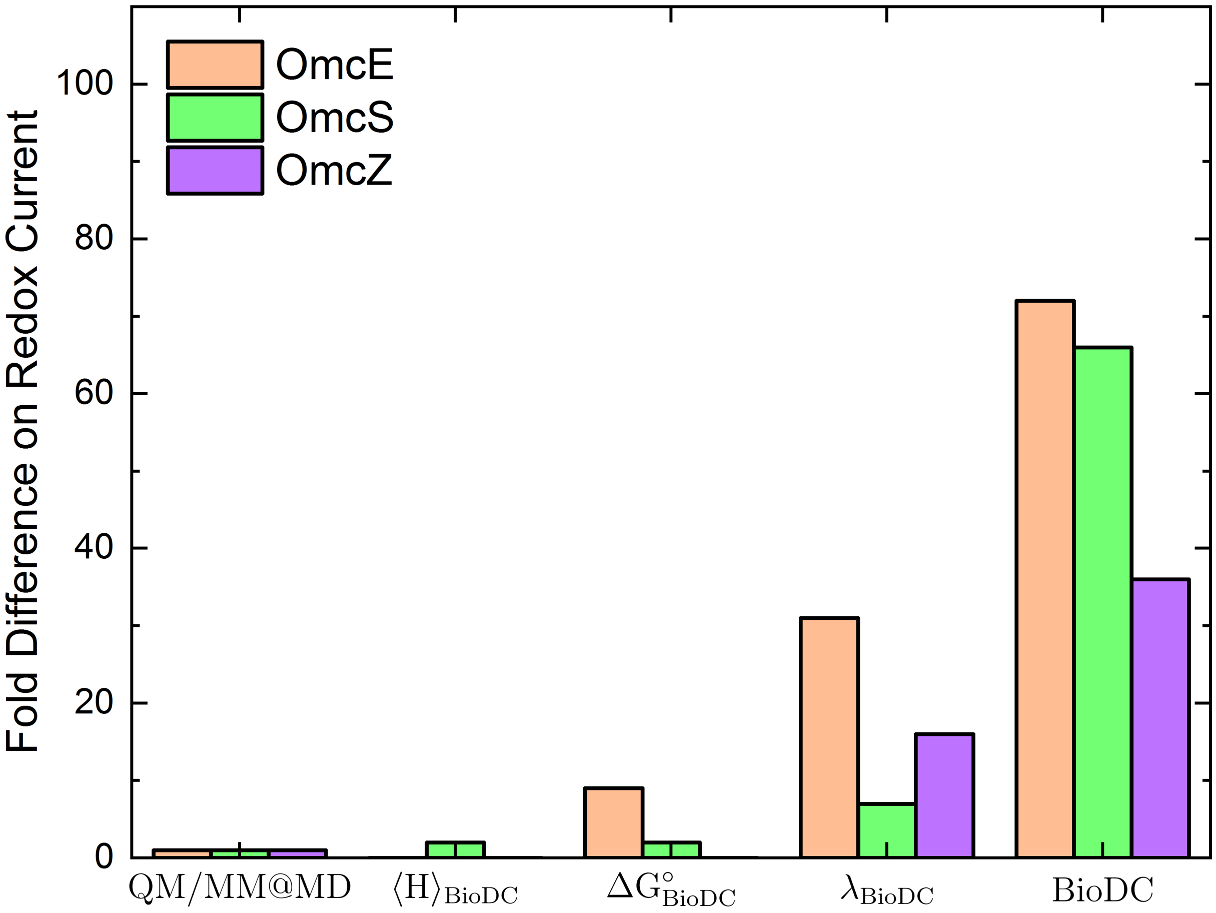


Figure S1. Fold differences relative to QM/MM@MD results as λ, ΔG°, and ⟨H⟩ computed with BioDC are, from left-to-right, used separately and then all-together to compute redox currents.
